# From Correlation to Communication: disentangling hidden factors from functional connectivity changes

**DOI:** 10.1101/2022.09.25.509384

**Authors:** Yuhua Yu, Caterina Gratton, Derek M. Smith

**Author notes:** Correspondence: Yuhua Yu.

## Abstract

While correlations in the BOLD fMRI signal are widely used to capture functional connectivity (FC) and its changes across contexts, its interpretation is often ambiguous. The entanglement of multiple factors including local coupling of two neighbors and non-local inputs from the rest of the network (affecting one or both regions) limits the scope of the conclusions that can be drawn from correlation measures alone. Here we present a method of estimating the contribution of non-local network input to FC changes across different contexts. To disentangle the effect of task-induced coupling change from the network input change, we propose a new metric, “communication change”, utilizing BOLD signal correlation and variance. With a combination of simulation and empirical analysis, we demonstrate that 1) input from the rest of the network accounts for a moderate but significant amount of task-induced FC change; 2) the proposed “communication change” is a promising candidate for tracking the local coupling in task context-induced change. Additionally, when compared to FC change across three different tasks, communication change can better discriminate specific task types. Taken together, this novel index of local coupling may have many applications in improving our understanding of local and widespread interactions across large-scale functional networks.

## Introduction

The brain is a dynamic system that integrates information in order to fulfill its functions. This system is characterized by both short and long range connections between regions which are organized into large-scale brain networks (Fox et al., 2005; Power et al., 2011; Yeo et al., 2011). The coupling in activity between different brain regions is referred to as functional connectivity (FC). FC is typically measured by indices of statistical dependence, such as correlation coefficients, coherence, or transfer entropy (Baccalá & Sameshima, 2001; Patel et al., 2006; Vicente et al., 2011). In human functional Magnetic Resonance Imaging (fMRI), the Pearson correlation derived from Blood-oxygen-level-dependent (BOLD) time series is the dominant means of quantifying FC and drawing inferences on neural coupling (Biswal et al., 1995; David et al., 2004; Fiecas et al., 2013; Mahadevan et al., 2021).

While functional connectivity patterns are largely stable over time within an individual (Gordon et al., 2017; Gratton et al., 2018; Laumann et al., 2015), there is evidence that task context induces subtle changes in FC (Arbabshirani et al., 2013; Cole et al., 2014; Gonzalez-Castillo & Bandettini, 2018; Gratton et al., 2016;2018; Krienen et al., 2014; Wu et al., 2021). Changes in FC between rest and task can help us understand how functional networks support human cognition (Allen et al., 2014; Chang & Glover, 2010; Kietzmann et al., 2019), but its interpretation remains controversial (Hutchison et al., 2013; Laumann & Snyder, 2021). Some authors have raised concerns about interpreting change in correlation as change in neural coupling (Behseta et al., 2009; Friston, 2011; Laumann et al., 2017). For example, Duff et al., (2018) suggested that many changes in BOLD correlation can be attributable to “additive signal change”, where a separate signal is added to one or both neighboring nodes. Additive signal changes include changes in noise levels or the amplitude of a common signal driving correlation between two nodes. By making an inference from the relationship between changes in variances and covariances, the authors distinguished additive signal changes from changes involving more complex combination of increases and decreases in the strength of existing signal components. A conjunction analysis incorporating variances and covariances was also proposed by Cole and colleagues (2016) to investigate the shared and unshared signals in BOLD data.

In the current project, we distinguish two types of factors driving the change in BOLD FC (Fig 1). The first type involves a change in the *local coupling* between two nodes. Local coupling can be altered by increased synaptic strength (e.g., Yao et al., 2007; Zucker & Regehr, 2002) in the cortical circuit. Local coupling can also be instantiated when the oscillation in a neuronal group is entrained by the oscillation from another group (Fries, 2005). Local coupling is closely related to the causal influence^1^ (Friston, 2011) between two regions, regardless of anatomical distance.

**Figure 1.**
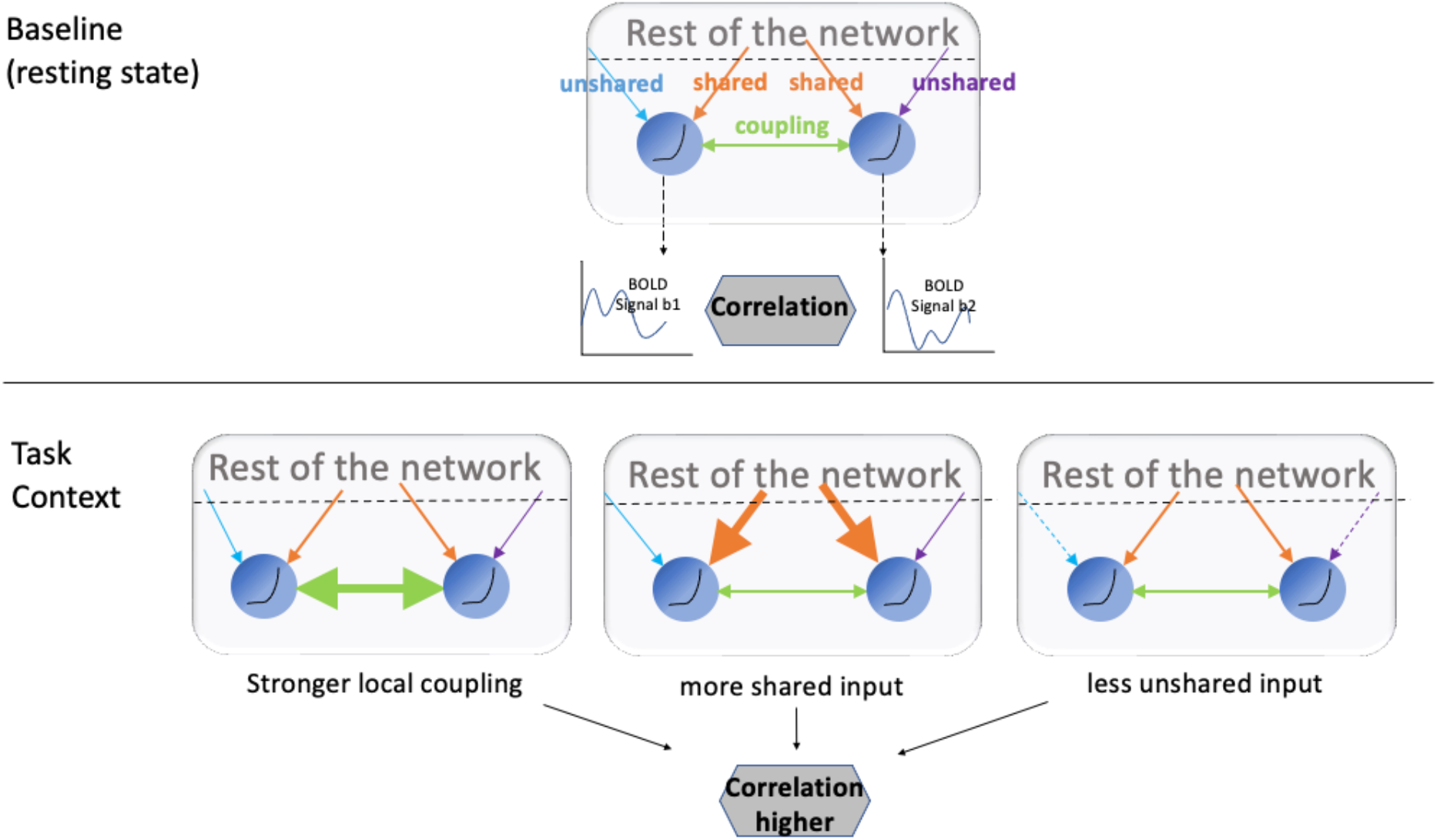
A schematic representation: different factors can drive functional connectivity across contexts. When the BOLD correlation between two regions increases, it can stem from stronger coupling between the two regions (direct synaptic input, etc), higher shared input from the rest of the network, or decreased unshared input that is idiosyncratic to one of the regions.

The second type of factor involves input from the rest of the network that modulates the FC of an edge even in the absence of causal interaction between two neighbors. We differentiate two subtypes here. The first subtype, *shared input*, is comprised of neural signals that simultaneously affect both neighbors. For example, increases in the correlation between visual regions during visual stimulation could be explained by shared activity induced by the same stimulus. There is also considerable evidence showing that the ascending arousal system plays a role in modulating functional network topography globally (Guedj et al., 2017; Shafiei et al., 2019; Shine, 2019; Shine et al., 2016;2018; Turchi et al., 2018). Projections from critical neuromodulatory sources like the dorsal raphe, locus coeruleus, ventral tegmental area, pedunculopontine nucleus, and the cholinergic basal forebrain are widespread thus making these regions potential sources of shared input (Aston-jones & Cohen, 2005; Mena-Segovia & Bolam, 2017; Morales & Margolis, 2017; Selden et al., 1998; Shine, 2019). The second subtype, *unshared input*, involves neural signals that are idiosyncratic to each node of a pair. For example, signals from different modalities (visual and auditory for example) can project to separate, but coupled regions in the higher order cortex (Sepulcre et al., 2012). Unshared input can also include local neural noises (Daunizeau et al., 2012) or measurement errors. Regardless of the mechanism, the two subtypes of input are exogenous to the causal interaction between two neighbors, and we collectively refer to them as (non-local) network input.

While neither network input, nor local coupling can be observed directly with fMRI, we aim to estimate their contributions to task-induced FC change by incorporating information regarding the temporal variance of the BOLD signal. Specifically, we focus on the FC changes under “task state” that occur over a block (multiple minute period). Like correlation, BOLD signal variance can change over contexts. For example, resting state BOLD data has been reported to have a higher variance than task BOLD data (Ito et al., 2020). However, for a pair of neighbors, the shared and unshared input variance can have opposing effects. Shared signals are expected to increase the correlation between the pair while unshared signals to attenuate the correlation. We aim to quantify this relationship in both simulation and real data by estimating the network input variance from BOLD signals.

Our simulation not only highlights the multiple factors modulating FC but also provides insight on a new measure, *communication change*, that may be used to track local coupling more closely than FC. Communication change utilizes a learned relationship between FC and input variances during a baseline (or resting state) to mitigate the impact of input fluctuation on task-induced FC change. However, unlike correlation or partial correlation (see below) that can be used as a “standalone” metric for functional connectivity, *communication change* is a method to disentangle factors underlying *FC changes* across task contexts ^2^.

Previous work has used partial correlation (Marrelec et al., 2006) to disentangle the impact of local coupling from the mutual dependencies on other brain regions. Partial correlation calculates the correlation between a pair of time series after the portion of their variance explained by all other observed time series is removed. Thus, partial correlation can capture a similar concept as *communication change*, but with a different modeling and computational approach. To evaluate how partial correlation is affected by the changes in local coupling and netwok input, we analyzed it in the same simulation framework.

When examining empirical data, we found that network input account for a significant but moderate amount of task-induced FC change. To estimate FC and input variances during task performance, we used BOLD signal residuals after removing evoked task effects following the ‘‘background connectivity’’ approach (Al-Aidroos et al., 2012; Fair et al., 2007). Depending on the task context, communication change sometimes suggests local coupling changes in the opposite direction as FC change would indicate. To gain insights into the informational content of *communication change* we investigated whether it can reliably discriminate task type.

## Results

### FC is sensitive to both local coupling and network input, while communication tracks local coupling more closely

We simulated two interacting neural regions with a two-dimensional Wilson-Cowan dynamic model (Wilson & Cowan, 1972.) to illustrate that 1) FC change is sensitive to local coupling as well as to network input; and 2) the impact of network input is dampened in the *communication change* measure where the contribution from the rest of the network, proxied by shared and unshared input variance (see *Methods-communication change*), is removed from FC changes. BOLD signals were computed from simulated neural activity via the Balloon-Windkessel model (Friston et al., 2003). FC and communication changes were derived from the simulated BOLD series. Throughout, we refer to the temporal correlation between nodes in this model as “FC”, in equivalence to BOLD correlation ^3^.

With all parameters (including the local coupling weight, and shared/unshared neural signal variances) fixed, we computed baseline FC, baseline input variance for each node, and regression coefficients of FC with respect to the input variance. These values will be used to compute *communication change* in the following scenarios: 1) after varying local coupling strength, 2) after varying shared neural signal variance; and 3) after varying unshared neural signal variance.

As expected, simulated FC is sensitive to both changes in local coupling and changes in input signal variance, confounding the interpretation of FC measures. Both stronger local coupling (Fig. 2A) and amplified shared signal (Fig. 2B) lead to higher FC, while amplified unshared signals (Fig. 2C) lowered the FC. In comparison, communication change shows a dampened relationship to shared and unshared signal variance while remaining sensitive to the local coupling (Fig. 2A-C). For example, with the current simulation setup, both FC and communication were equally sensitive to local coupling. However, FC was 5 times more sensitive to shared signal (the slope in Fig. 2B) and 2.5 times more sensitive to unshared signal (the slope in Fig. 2C) relative to communication.

**Figure 2.**
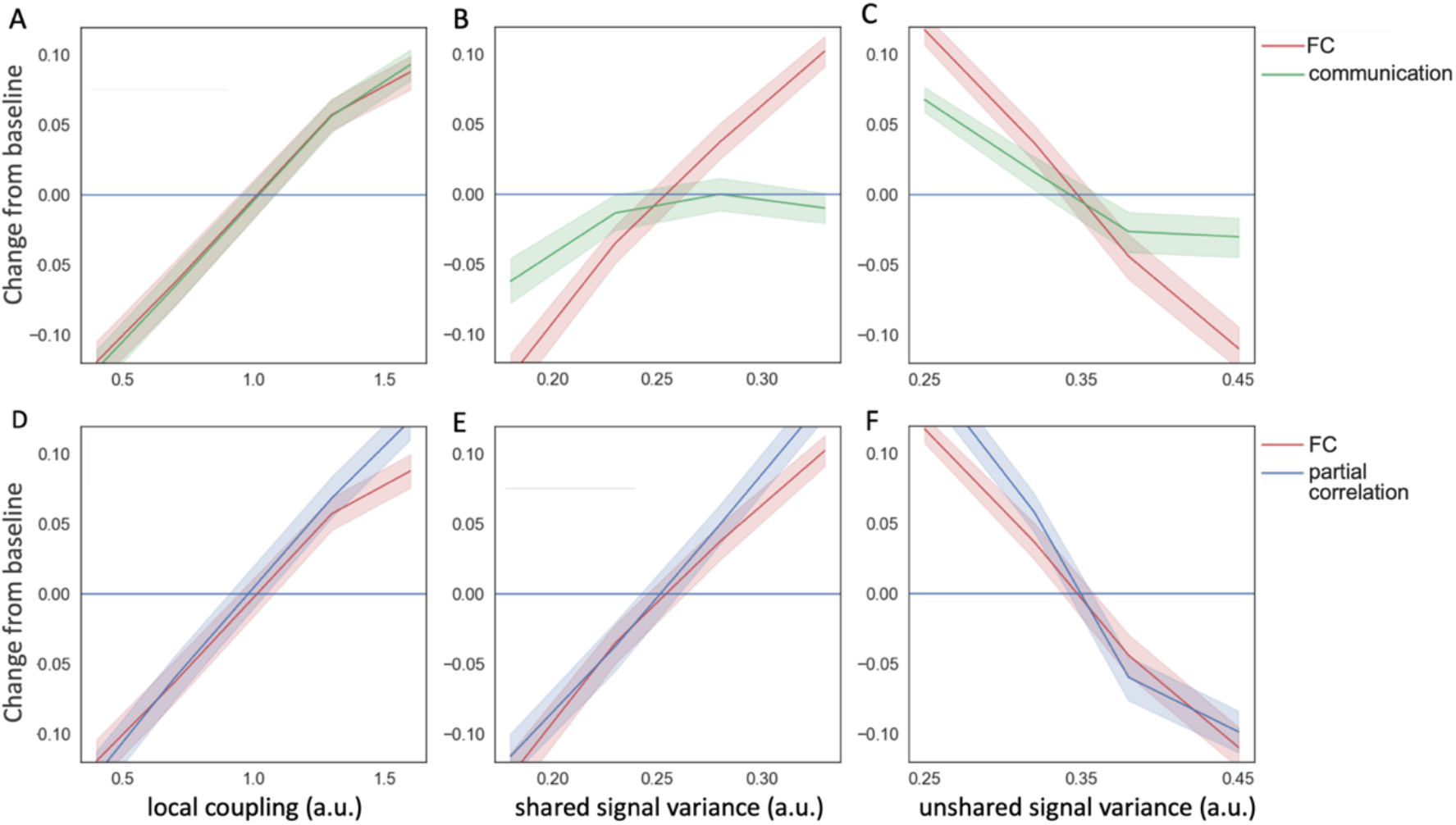
The sensitivity of FC, communication and partial correlation with respect to model parameters (A) Both FC and communication increase with the coupling strength (W). (B) FC (but not communication) increases sharply with increases in shared signal variance. (C) FC decreases with unshared signal variance while the effect is dampened for communication. (D-F) FC and partial correlation in the same simulated scenarios display similar behavior to one another.

Given that partial correlation is often used to account for the mutual dependence on other brain regions, we computed partial correlation in the same three scenarios (see *Methods*). We asked: does partial correlation have less dependence on input signal variance than FC? In the current simulation framework, partial correlation had similar sensitivity as FC with respect to changes in local coupling and input signals (Fig. D-F). Therefore, it cannot discriminate among those factors.

This analysis suggests that communication, compared to correlation and partial correlation, more closely tracks changes in local coupling relative to changes in input variance. Communication diminishes the impact of network input by using the statistical relationship between FC and variance in a baseline state. Notably, this is an imperfect proxy because input signals are unobservable in real data. Therefore, communication is not guaranteed to have zero sensitivity to shared and unshared input variances, but their impact is considerably dampened as observed from the simulation. This property allows us to infer the change in local coupling from real data when neither the coupling nor input signals are directly observable.

### FC is related to shared and unshared input variance in real fMRI data

To compare communication and FC in empirical data, we used Midnight Scan Club dataset (Gordon et al., 2017), a precision fMRI dataset with 10 sessions of fMRI rest and task data from 9 participants. See *Methods* for BOLD data acquisition, preprocessing, FC, and variance calculation for resting state and task data. We computed BOLD FC, shared input variance, and unshared input variance for each of the 10 resting state scans. We fit a linear model for each edge with the FC as the dependent variable and shared/unshared input variance as independent variables. As expected, FC increased with shared input variance and decreased with unshared input variance in the MSC data. Figure 3 shows the distribution of the regression coefficients of all participants. Network input fluctuations jointly explained 24.7±0.1% (adj. R-square) of the FC variability across resting state sessions. Results from individual participants all show a similar pattern and are included in Supplemental Fig. S1, indicating robustness of the results.

**Figure 3.**
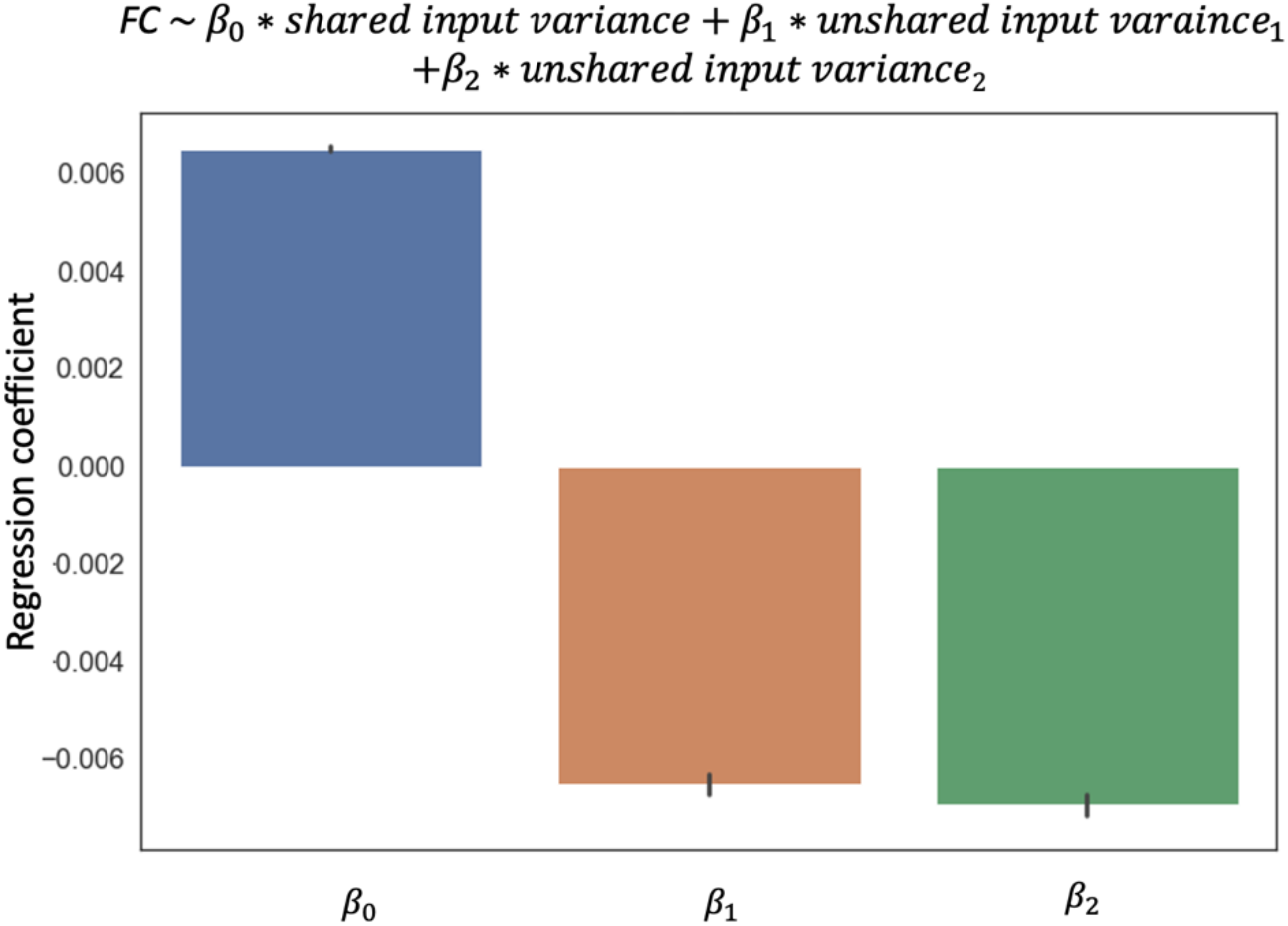
Distribution of regression coefficients of FC against the unshared and unshared input variance in the Midnight Scan Club. The regression is run for each edge across resting state scans for each participant. Error bars show a standard error across edges. Data from all participants and edges are plotted here. Results for individual participants are shown in Supplemental Fig. S1.

Thus, the empirical relationship between FC and network input is consistent with what we observed in the simulation: FC increases with the shared input variance and decreases with unshared variance idiosyncratic to one of the nodes. We next use the regression coefficients learned from resting state to estimate the contribution of input variance in FC during task-induced contexts.

### Task-induced FC changes are related to changes in network inputs

Next, we asked whether task-induced FC change can be predicted by network inputs, estimated from variance changes. We computed the predicted FC changes during task performance from the observed input variance change under (Eqn.1). There was a significant correlation between the predicted and actual FC changes across tasks for the majority of the edges (Fig. 4 for the aggregated group distribution of the correlations, mean is 0.152±0.001); individual participants are shown in Fig. S2). This indicates that input fluctuation explains a small but significant proportion of task-evoked FC changes. The prediction error, i.e., *communication change* in our definition, is the part of the FC change not explained by input fluctuation.

**Figure 4.**
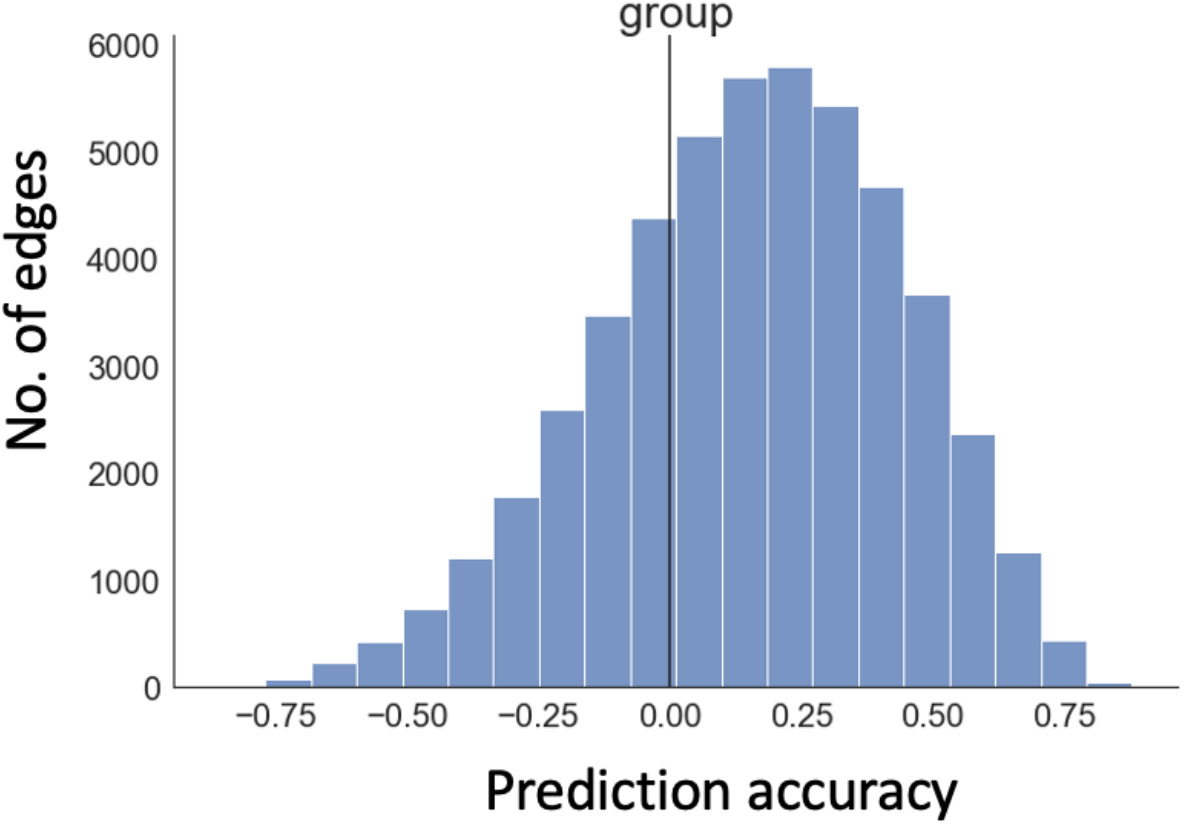
Ability of changes in input variance to predict FC changes during tasks in real fMRI data from the Midnight Scan Club. For each edge, accuracy was measured using the correlation coefficients between actual FC changes and those predicted by task-induced input variance changes across 30 task conditions (10 sessions for memory tasks, 10 for motor task, and 10 for mixed task). Data plotted here is based on the group average; individual participant relationships are shown in Fig. S2.

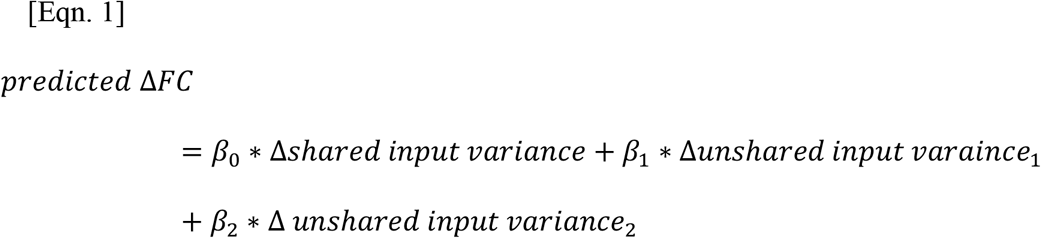

*where β’s are computed from resting state data (Fig. 3) and input variance changes are measured from the comparison in variances between task and rest*.

### Communication change differs from FC change and shows improved sensitivity to task information

Since the previous simulations suggest that *communication* tracks local coupling while dampening the impact of input fluctuation, we next directly compare FC and communication measures in real data. We compared the task-induced changes in each measure, grouping edges according to a widely used large-scale network designation (Gordon et al., 2016). Communication and FC changes during tasks differed in many networks and these differences were modulated by the task type (Fig. 5). Sometimes communication was higher magnitude than FC, and other times it was lower; on occasion it even showed relationships in the reverse direction. For example, when participants performed the memory task (Fig. 5A), FC decreased on average within the ventral attention network, while communication (calculated from FC after removing the contribution from input variance, see *Methods* eqn. 2) increased. This highlights the danger of interpreting an FC decrease directly as reduced inter-regional interaction.

**Figure 5.**
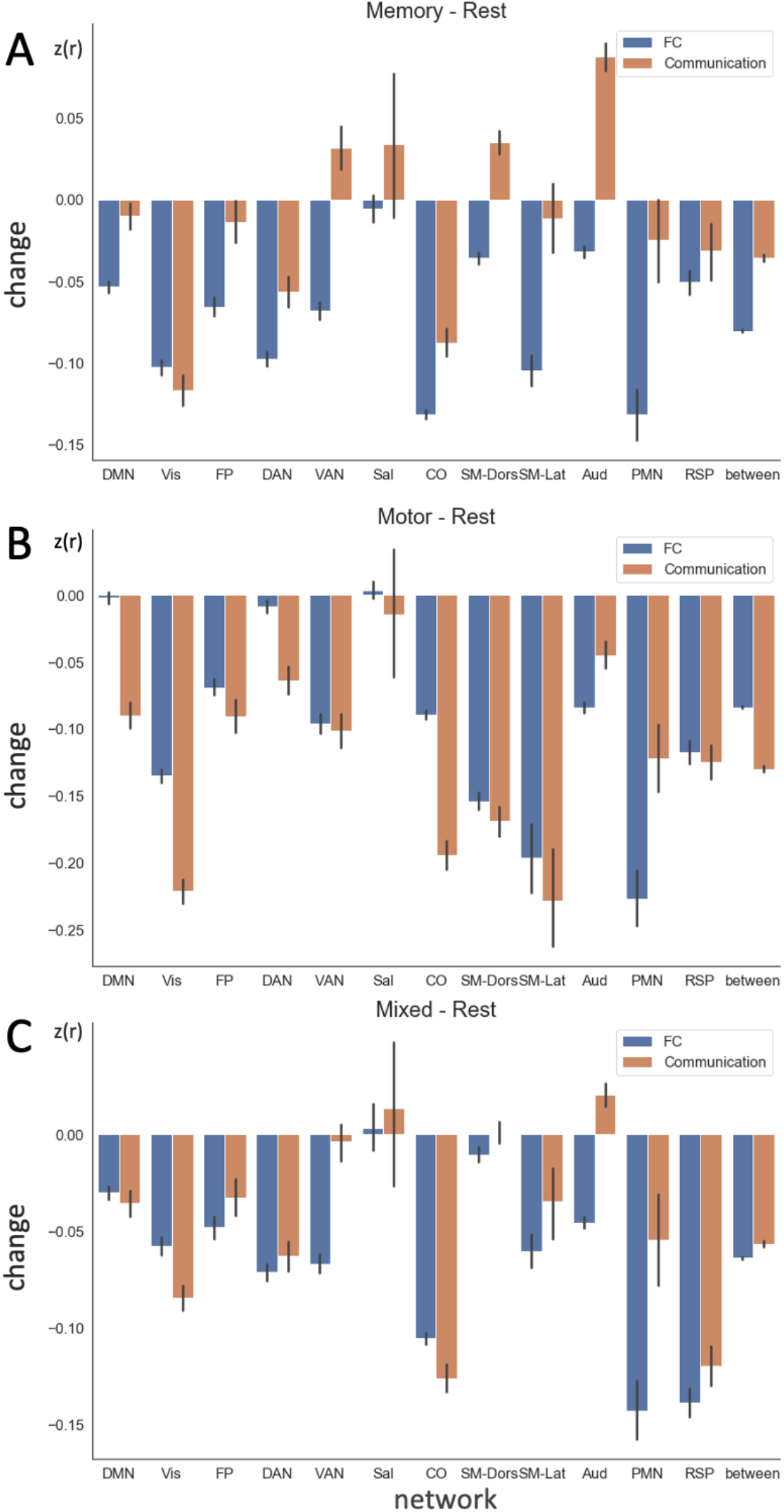
Group averaged, task-induced changes in FC and communication for all edges grouped by network affiliation during (A) memory task, (B) motor task, and (C) mixed task (dot coherence and semantic) relative to rest in the MSC. Error bar indicates the standard error across edges. Data from individual participants are shown in Fig. S3. DMN: default mode; Vis: Visual; FP: frontoparietal; DAN: dorsal attention; VAN: ventral attention; Sal: salience; CO: cinguloopercular; SM-Dors: dorsal somatomotor; SM-Lat: lateral somatomotor; Aud: auditory; PMN: parietal memory; RSP: retrosplenial; between: between network connectivity.

To probe the informational content in the communication measure, we ask to what extent it can discriminate among tasks with distinct task demands. To this end, we computed the intraclass correlation coefficient (ICC, see *Methods*) between task types for each edge across all scan sessions (4 task types including the resting state with 10 sessions each). ICC measures the percentage of variance accounted for by task types among all changes observed for the same edge (Bartko, 1976; McGraw & Wong, 1996). If communication change contains task information, ICC^4^ is expected to be closer to 1 (rather than to 0).

Across all edges and all participants, the average Communication ICC = 0.699±0.001, with 71% of the edges showing a a *p*-value less than 0.05 (F-test), indicating that most edges contain reliable information about task type (Fig. 6). For comparison, we computed the ICC for FC changes using the same methodology. Similar to communication, FC changes reliably represent task information (ICC = 0.462±0.001; Fig. 6). However, the ICC based on the communication measure is significantly higher than the ICC of FC (*z* = 140, *p*<.00001), suggesting that communication change increases the discrimination of task type.

**Figure 6.**
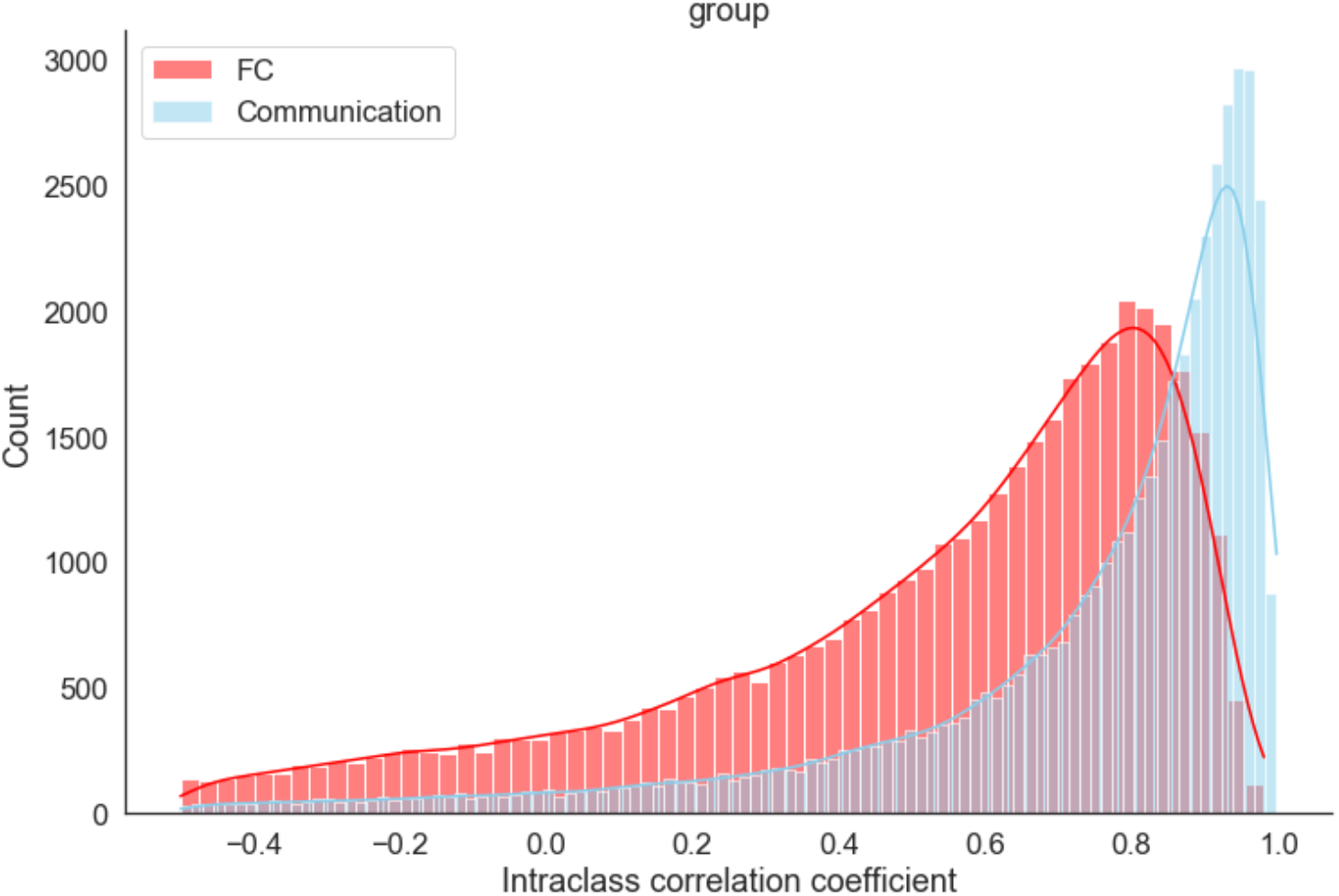
Distribution of Intraclass correlation coefficients (ICC), across all edges and participants, for discriminating among task conditions. The ICC between task types was computed for each edge across 40 scans (10 each for rest, memory, mixed, and motor). Data from individual participants are shown in S4.

The same pattern was observed consistently for all participants (Fig. S4). The difference between FC and communication in terms of task discrimination is not solely driven by the distinction between the resting state and *any* task state. ICC among the three task states exhibits a similar pattern (although the difference is narrower, *z* = 77, *p*<.00001, see Fig. S5) indicating that communication change contains more task specific information relative to FC.

In summary, communication changes during tasks exhibit significant differences from FC in various networks of the brain. Importantly, communication contains more informational content to discriminate specific task types, compared to FC.

## Discussion

While BOLD correlation is widely used to capture both static functional connectivity and its changes across contexts, the interpretation of this measure is often ambiguous. The entanglement of multiple latent factors including local coupling of two neighbors and network inputs from the rest of the network limits the scope of the conclusions that can be drawn from correlation measures alone. Here we developed a method to compute the shared and unshared input variance to an edge and quantify their impact on task-induced FC changes. We further proposed a metric, termed “communication change”, that dampens the impact of input signals and thereby provides a promising candidate for more closely tracking local coupling changes.

We first analyzed local and network impact on FC by conducting a simulation based on a well-established model of neural dynamics (Wilson & Cowan, 1972). The model suggests that FC can be driven by multiple factors. Increased local coupling as well as increased variance of shared inputs result in higher FC, while increases in the variance of unshared (idiosyncratic) input has the opposite effect. Although the conclusion seems intuitive, explicit modeling of the three latent factors leads to a novel metric, communication, that dampens the impact of both shared and unshared inputs using only observable, statistical relationships. Our simulation shows that communication change is more likely to track local coupling.

Interestingly, partial correlation, a metric often used to disentangle the impact of local coupling from the mutual dependencies on other brain regions (Marrelec et al., 2006; Smith, 2012), shows similar sensitivity as FC change in the current simulation framework. Partial correlation removes the contemporaneous signal dependence using a linear residualization. However, as the BOLD signal exhibits temporal nonlinearity, the transformation from neural activity to BOLD signal (via Ballon-Windkessel model) is state and path dependent (Buxton et al., 2004; Friston et al., 2000). It is possible, therefore, that the correlation between residualized signals is still affected by the network input through their “history”. We are not aware of other simulative studies illustrating the (lack of) efficacy of partial correlation, except for (Cole et al., 2016), where the authors also reported that partial correlation retains limitations of the FC measure. In comparison, to compute communication change, we first aggregated inputs from the rest of the network into three terms, shared variance and two unshared variances idiosyncratic to each node. Communication then removes the input contribution via a “learned” statistical relationship between FC and variance at baseline (here operationalized as the average resting-state signal). Therefore, the method makes few assumptions about the underlying structure of the input signals.

With empirical data, we computed the sensitivity of FC with respect to shared and unshared input during resting state scans. As expected, FC in real BOLD data increased with the variance of shared input and decreased with unshared input. We then analyzed task-induced FC changes using the coefficients learned from the resting state. Input fluctuation explained a significant, but moderate, amount of FC changes during task performance. After removing the input related contributions, the remaining FC change, i.e., communication change, often had the opposite sign of the original FC change in various tasks and brain networks. This highlights the ambiguity of interpreting task-induced FC change by itself. If the correlation of an edge increases, for example, it can be driven by an increase in shared input from the rest of the network, a decrease in an idiosyncratic input affecting only one of the nodes, an enhanced neural coupling between the nodes, or any combination of the above factors.

Although we cannot directly observe how communication differentiates mechanisms behind FC changes in empirical data, evidence suggests that it shows better discrimination across the task types relative to the standard FC measure. Both simulation and empirical analysis suggest that communication may be an improved measure (from FC) for tracking changes in causal interactions between regions across task contexts. Such changes can occur at the synaptic level due to short-term plasticity (Krystal et al., 2003; Yao et al., 2007; Zucker & Regehr, 2002), such as an elevated dopamine tone in the same cortical circuit (Vijayraghavan et al., 2007), and via increased coherence or phase locking (Fries, 2005).

While FC is strongly related to shared and unshared network inputs, communication shows a diminished relationship to these latent signals. Shared input can arise from signals with widespread influence stemming from subcortical projections (Raut et al., 2021; Shine, 2019), while unshared (idiosyncratic) input can come from, for example, unique inputs from different modalities (visual or somatosensory) that only project to selected regions. The communication metric dampens the impact of network input changes using a statistical relationship and remains agnostic to the underlying biophysiological mechanism. This allows contextual factors to guide the interpretation of the difference between FC and communication across a variety of states.

### Using BOLD variance to augment the interpretation of FC

Other researchers have also explored the idea of incorporating variance (or covariance) to disambiguate the interpretation of FC (Duff et al., 2018; Cole et al., 2016). For example, Duff and colleagues (2018) combined correlation and variance to classify FC changes into cases attributable to “additive signal change” (a type of “shared or unshared input”) and changes arising from modifying existing signals. While the current analysis echoes the importance of augmenting FC with variance/covariance, we propose a framework to quantitatively decompose FC changes into contributions from different underlying mechanisms.

Communication shows higher intraclass correlation across tasks compared to FC, suggesting that the proposed measure better discriminates task types. This suggests that local coupling is modulated by specific task demands, while overall signal variance contains less task-specific information. Previous work has shown a simultaneous task-induced reduction in FC and variance (Ito et al., 2020). It is possible that task performance, relative to the resting state, reduces synchronized, spontaneous activity that affects a large region of the brain and leads to an FC reduction in a pattern that is similar across task types. By extracting the “excess” FC change after accounting for the variance-related effects, we can gain more insight into how specific task demand modulates the brain network.

Furthermore, the current framework provides a way to decompose exogenous network inputs into shared and unshared variance. One powerful approach may be to use several of these metrics in combination. For example, an increase in communication implies stronger local coupling. If a decrease in FC is simultaneously observed, we can ask whether the discrepancy is due to a reduction of shared signal or an increase of idiosyncratic inputs by examining changes in shared and unshared variance simultaneously. Therefore, examining communication together with FC may allow researchers to draw inferences regarding how a particular region adjusts its interactions with its neighbors across a range of different contexts, providing an improved handle on whether local or shared inputs are at work in any given scenario.

### Limitations and future directions

One pitfall of our approach is that the interpretation of our simulation results is dependent on the validity of our computational model. The communication change metric was inspired by a simplified neural dynamic simulation. Although Wilson-Cowan dynamics have been widely used and have had a major impact on computational neuroscience, it is a simplification of the underlying biological process (Chow & Karimipanah, 2020). Many extensions and generalizations have been proposed since its initial introduction in 1972. To highlight the basic interaction between several statistical measures of interest, we adopted the simplest form of the model in a two interacting nodes set-up. By doing this, we imposed the assumption that any effect from the rest of the system could be represented by Gaussian signals. Furthermore, we fixed the mean of the input signals and focus on the impact of their variance/amplitude. Because the neural dynamics involve sigmoid activation functions, the shift in the average level of the input signal would have a non-linear impact on the resulting fMRI BOLD signals (Ito et al., 2020). Future work will explore how the average activity level interacts with the relationship between FC and input variances discussed in this article.

When it comes to empirical data, a key premise of our analysis is based on evidence that fMRI signals are indirectly related to underlying neural population activity. Such a view is widely supported in the literature (Logothetis, et al., 2001; Ma et al., 2016), but certain MRI artifacts can ambiguate the results (Friston et al., 1996). We employed multiple artifact mitigation measures (scrubbing, motion regression, global signal regression, white matter signal regression, and ventricle signal regression) to minimize the impact of artifactual factors.

Moreover, the BOLD signal analyses utilized 9 highly sampled subjects from the MSC dataset (Gordon et al., 2016). This is a small sample, but there is evidence that results from MSC data extend to other independent datasets (Gordon et al., 2018). Importantly, the single subject results presented here are consistently observed in all individuals, demonstrating a certain degree of robustness. However, communication change should be examined in other data sets in future work, especially with task conditions that impose varying degrees of cognitive demand and a larger number of subjects.

There are unanswered questions that are beyond the scope of the current investigation that pertains to communication change. Task-induced communication increases in some regions (e.g., visual network, dorsal attention network) and the magnitude of the changes seems to be modulated by the type of task. In addition, the spatial pattern of communication exhibits significant individual differences (Fig. S3), consistent with findings from previous work (Gratton et al., 2018; Porter et al., 2022). An intriguing question for future research is how differential communication is related to task performance.

Finally, task discrimination is only one possible venue to explore the informational content in communication and its advantage over FC. Research directly modifying potentially widespread signals such as arousal (shared input), signals that might be expected to be more local such as selective stimulus-response features (unshared input), or manipulation at the local synaptic level (coupling) may help to validate the interpretation of communication change.

### Conclusions

FC, computed as a correlation between paired BOLD signals, is not only determined by local coupling between pairs of regions but also by inputs from the rest of the network, among other factors. Utilizing the statistical relationship between FC and input variance, we developed a method to extract a component (“communication change”) from FC change with a dampened influence of network input. Examining changes in communication, along with FC, allows us to draw inferences regarding how the interaction between a particular region with its neighbors changes across different contexts.

## Materials and Methods

We used a combination of numerical simulations and empirical analysis to examine the effects of local interactions and input fluctuations on BOLD correlation and variance. These results led us to develop a new measure, communication change, that more closely tracks the change in coupling between two neural regions compared to the standard FC measure (BOLD correlation) across task contexts. Below, we first describe our modeling approach, then the derivation of the communication metric, and finally we include a description of a dataset and analyses used to empirically examine differences in communication change and correlation.

### Compute communication change in both simulated and real data

We developed a novel metric to better capture the inter-regional coupling shift in task-induced FC change, which we termed “*communication change”*. To compute the communication change of an edge, we first need to estimate the shared and unshared network input variance.

<u>Shared/unshared input variance</u> is computed from the variance of the BOLD signals from common and idiosyncratic neighbors. Specifically, assume *node1* and *node2* are the nodes of the edge under consideration. *N*_*s*_ is the set of neighboring nodes that connect to both *node1* and *node2. N*_*1*_ is the set of nodes that only connect to *node1*, and *N*_*2*_ is the set of nodes that only connect to *node2*. I.e., *Ns, N*_*1*_, and *N*_*2*_ are mutually exclusive. Shared input variance is the total variance of the BOLD signals from *N*_*s*,_ computed by the trace of the covariance matrix. We residualized all the signals in *N*_*1*_ against all signals in *N*_*s*_. *Unshared input varaince*_1_ is the total variance of the residualized signals in *N*_*1*._ Likewise, *unshared input varaince*_2_ is the total variance of the residualized signals in *N*_*2*_ against *N*_*s*._

*<u>Communication change</u>* is computed with the following equation:

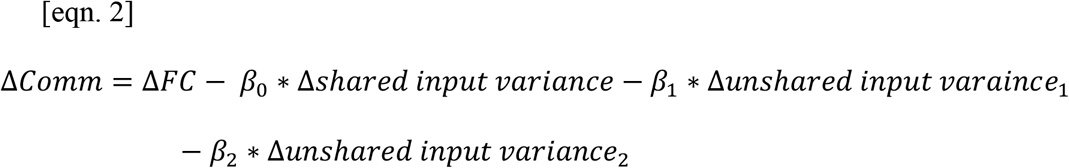

Where all the changes are based on comparisons of task-rest, or for the simulation, a scenario with parameter shock – baseline.

The coefficients *β*_0,_ *β*_1,_ *β*_2_ are then computed from resting state (or the baseline simulation scenario) using linear regression:

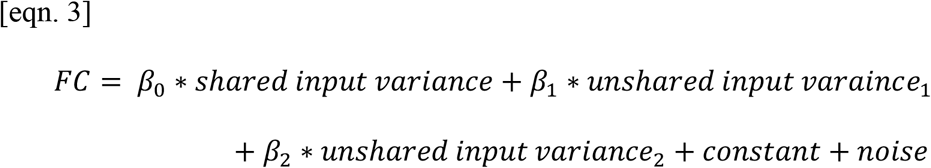

All edges and neighbors in empirical data (including rest and task) are defined by 5% sparsity based on the average resting state correlation. We reevaluate the key findings using 10% sparsity and reached similar conclusions (Supplement S6).

### Numerical simulation of BOLD signals from two interacting neural regions

We examined the effects of local coupling and network inputs on BOLD correlation with a numerical simulation. A two-dimensional (two node) Wilson-Cowan type (Wilson & Cowan, 1972) dynamic was used to describe the activity of two interacting neural regions:

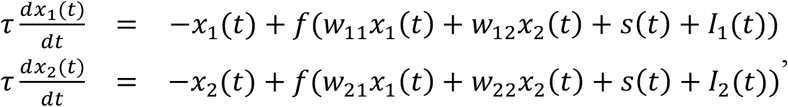

where *x_i(t)* represents the excitatory level of a neural group.

The matrix 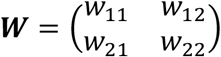 describes the information exchange within and between regions. In particular, the off-diagonal terms, *w*_21, *w*_12, determine the strength of inter-region coupling. For the baseline scenario, the matrix was set to 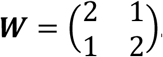. The diagonal entries were fixed throughout the simulation. The parameter values (and those in the following) are adapted from (Ito et al., 2020) and adjusted to obtain a realistic baseline correlation.

Shared input *s(t)* is a Gaussian signal with shared signal variance *V*_*sh*,_ which was set to *V*_*sh*_=0.25 in the baseline scenario. *V*_*sh*_ indexes the magnitude of the common exogenous input from the rest of the brain network to both regions.

*I*_1 (*t*), *I*_2 (*t*) are two independent Gaussian signals representing idiosyncratic signals with unshared signal variance parameter *V*_*un*_, which was set to 0.35 in the baseline. *V*_*un*_ indexes the magnitude of the idiosyncratic input to one of the neighbors.

*f* is the sigmoid input-output activation defined as *f*(*x*) = 1/(1 + *e*^(−*k* * *x*)), where *k* is fixed at 0.5 throughout the simulation.

Neural activities simulated from the above equations were transformed to BOLD signals using the Balloon-Windkessel model (Friston et al., 2003). The transformation assumes normalized deoxyhemoglobin content, normalized blood inflow, resting oxygen extraction fraction, and normalized blood volume. State equations and parameters were taken from previous work (Friston et al., 2003). Correlation (FC), partial correlation, and communication change were then computed on the simulated BOLD signals as specified below. Each simulation (1 run) generated a pair of BOLD timeseries with 2000 steps each.

For the <u>baseline simulation,</u> we ran the scenario with fixed parameters 500 times (500 runs). We obtained the average FC, average shared/unshared input variance, and the regression coefficients as in Eqn. 3. The baseline simulation mimics the resting state as in real data.

Next, we computed the sensitivity of correlation, communication, and partial correlation with respect to the input signal variance and local coupling parameters. To this end, we carried out the same simulation as above, but now independently varied the shared signal variance *V*_*sh*_, unshared signal variance *V*_*un*_ and the coupling parameter *w*_12, (and symmetrically *w*_21). We used inputs for *V*_*sh*_ ranging from 0.15 to 0.35, *V*_*un*_ ranging from 0.25 to 0.45, and *w*_12, from 0.4 to 1.6 (centered around baseline 1). The range was chosen such that the parameters had comparable impact on FC. For each scenario, the model is run 500 times to obtain the mean and standard deviation of all quantities of interest.

<u>Correlation (FC)</u> was then computed between a pair of simulated BOLD timeseries.

<u>Communication change</u> was computed as described above. All the variances required in the formula were computed from the simulated BOLD signals. To simulate the input BOLD signals, we applied the Balloon-Windkessel model to transform the shared signal *s*(*t*) and the idiosyncratic signals *I*_1_(*t*), *I*_2_(*t*).

<u>Partial correlation</u> was computed as the correlation between a pair of residualized series, after removing the common variance from the rest of the network. In the current simulation framework, inputs from the rest of the network are represented by the shared/unshared signals. Therefore, we residualized the BOLD signals derived from *x*_1_(*t*), *x*_1_(*t*), against the input BOLD signals derived from *I*_1_(*t*), *I*_2_(*t*) *and s*(*t*), and computed the correlation between the residualized signals.

### MSC dataset

In the second portion of our investigation, we examined the relationship between FC, variance, and communication in an empirical dataset. We used the Midnight Scan Club (MSC), a precision fMRI dataset (Gordon et al., 2017) that contains large quantities of both task and resting state data was selected, as large amounts of data are required to achieve highly reliable FC measures (Gordon et al., 2017; Laumann et al., 2015). The dataset includes data from 10 participants who each participated in 10 separate fMRI sessions. During each session, each participant completed 3 different tasks (a motor task, a memory task, and a mixed dot coherence/verbal discrimination task) and rest. This dataset is described in detail elsewhere (see Gordon et al., 2017), but we will review parameters relevant to this study below. In addition, a dataset of 120 healthy adults (WashU-120) was used as a reference set for the network definition and the calculation of graph theoretic metrics (Power et al., 2011; Power et al., 2013). This dataset has been described in detail in Power et al (2013).

### Task design

Each **resting-state** session was 30 min, during which participants were asked to lie still while fixating on a white cross presented against a black background. Each **motor task** session included two runs (7.8 min. total) of a blocked motor task adapted from the Human Connectome Project (Barch et al., 2013). In each block (15.4 s in duration), participants were cued to move either their left or right hand, left or right foot, or tongue. Each run included two blocks of each type of movement, as well as three fixation blocks (15.4 s). Each **mixed task** session included two runs (14.2 min. total) that each included four blocks, two of the semantic tasks and two of the coherence tasks. Both tasks had a mixed block/event-related design, modeled after the tasks in (Dubis et al., 2016). During the semantic task 30 individual trials were presented, consisting of words presented for 0.5 s with jittered 1.7-8.3 s intervals. Participants were asked to respond whether the words were nouns or verbs (50% nouns and 50% verbs were included). Forty-four second fixation periods separated blocks. During the coherence task (also 30 trials), individual trials consisted of arrays of Glass patterns (Glass, 1969) that were varied in how concentrically they were arranged (50% or 0% coherence to a concentric arrangement, displayed with equal frequency). Participants were asked to respond whether dots were arranged concentrically or randomly. Each **memory task** session included 3 runs of an event-related categorization task with implicit repetitions, with a separate run per stimulus type (face, scenes, and words). Within each run, participants viewed 24 images, repeated 3 times and made binary decisions.

### MRI acquisition parameters

High-resolution T1-weighted (224 slices, 0.8 mm3 isotropic resolution, TE = 3.74ms, TR = 2400ms, TI = 1000ms, flip angle = 8 degrees), T2-weighted (224 slices, 0.8 mm3 isotropic resolution, TE = 479ms, TR = 3200ms) both with 0.8 isotropic resolution, and resting state BOLD data were collected on a Siemens 3T Magnetom Tim Trio with a 12-channel head coil (Gordon et al., 2017). Functional scans were collected with a gradient-echo EPI sequence, isotropic 4mm^3^ voxels, TE of 27ms, and TR of 2200ms (Gordon et al., 2017). The MSC dataset is considered a deep (precision) dataset with 5 hours of resting state data per subject in 30 min. blocks over 10 separate sessions (Gordon et al., 2017).

### General Preprocessing

The T1-weighted images were processed via automatic segmentation of the gray matter, white matter, and ventricles in Freesurfer 5.3 (Fischl et al., 2002). The default recon-all command in Freesurfer was then applied to produce the anatomical surface for each subject (Dale et al., 1999). These surfaces were manually edited to improve the quality of the registration and the surfaces were registered to the fs_LR_32k surface space via the procedure outlined in Glasser et al., (2013). Functional data preprocessing began with field inhomogeneity distortion correction using the mean field map, which was applied to all sessions. Slice timing correction was implemented using sinc interpolation to account for temporal misalignment in slice acquisition time. This step was followed by motion correction which was conducted within and across BOLD runs (aligned to the first frame of the first run) through a rigid body transformation. Then whole-brain intensity values across each BOLD run were normalized to achieve a mode value of 1000 (Miezin et al., 2000). Functional BOLD data was first registered to a T2-weighted image and then to the T1. An affine transformation was employed for registration. The T1-weighted image was aligned to a template atlas (Lancaster et al., 1995) conforming to Talairach stereotactic atlas space (Talairach and Tournoux, 1988) using an affine transformation. All computed transformations and re-sampling to 3 mm isotropic voxels were simultaneously applied at the end of these steps.

### Functional Connectivity Pre-processing

Steps were taken to attenuate the impact of artifacts on BOLD time series. The effect of nuisance signals was mitigated through regression of average signal from the white matter, ventricles, whole brain (global signal), motion parameters, and derivatives and expansion terms of the motion parameters (Friston et al., 1998; Power et al., 2014). Removal of frames with FD > 0.2 mm, in addition to sequences containing less than 5 contiguous low motion frames, the first 30 seconds of each run, and runs with < 50 low motion frames was performed as an additional safeguard against the impact of motion (Power et al., 2014). A special filtering procedure was applied to two MSC subjects (MSC03 and MSC10) with respiratory contamination in their motion parameters. This procedure involved applying a low-pass filtered at 0.1 Hz to reduce the effects of respiratory artifacts on motion estimates stemming from the short-TR multi-band acquisition, all before censoring high-motion frames (Fair et al., 2020; Gratton et al., 2018; Laumann et al., 2017). Then a filtered FD threshold of 0.1 mm was applied to censor frames. In all cases, flagged head motion frames were removed and the time points were replaced with interpolated data using a power-spectral matched approach (Power et al., 2014), and this was followed by the application of a bandpass filter (0.009 Hz-0.08 Hz). Previous research has shown that ∼30 min. of low motion data is necessary to achieve high reliability of FC (Gordon et al., 2017). We therefore removed a participant (MSC08) that exhibited high degree of motion (Gordon et al., 2017). The excluded participant also reported a high level of drowsiness (Gordon et al., 2017; Laumann et al., 2017; Tagliazucchi & Laufs, 2014).

The processed BOLD data were mapped to each individual’s native midthickness surface via the ribbon-constrained sampling procedure (Marcus et al., 2013) Then, the mapped data were registered to the fsaverage surface in one step using the deformation map generated from the ribbon-constrained sampling procedure described in Glasser et al., (2013). This was followed by spatial smoothing conducted via a geodesic Gaussian smoothing kernel to the surface registered data (FWHM = 6 mm, sigma = 2.55) (Gordon et al., 2016; Marcus et al., 2011). Next, temporally interpolated frames were removed (prior to functional connectivity analysis).

### Regions, Networks

We examined a previously published set of 333 brain parcels associated with 12 networks Somatomotor Dorsal, Somatomotor Lateral, Visual, Auditory, Cinguloopercular, Frontoparietal, Dorsal Attention, Ventral Attention, Salience, Default, Parietal Memory, and Retrosplenial (Gordon et al., 2016).

### Task Functional Connectivity

In the case of the task scans, task related activation was captured with a general linear model (GLM) using in-house software written in IDL (Research Systems, Inc.), as described in (Gratton et al., 2018), using a finite impulse response approach for trial-level regressors (Ollinger et al., 2001). Functional connectivity was then measured for the task scans following the “background connectivity” method (Al-Aidroos et al., 2012; Fair et al., 2007) with the residuals from the GLM analysis being used for BOLD time-series correlations. The residuals were subjected to the aforementioned connectivity processing pipeline prior to functional connectivity calculation. There are multiple methods to measure task FC, that here we focus on background connectivity because of its simplicity and ability to track task changes at the resolution we are interested in, and that it is well suited for comparison with a resting baseline.

### Intra-class correlation coefficients to quantify task discrimination

We used the Python package Pingouin (Vallat, 2018) to evaluate how much variability in FC and communication across scans is attributable to task type. Specifically, we used the function *intraclass_corr*, where the “targets” are the task types (rest, mixed, memory, motor), raters are the session index (1 – 10). We used “ICC1k” output indicating that the target (task type) is rated by different raters (the session) and the raters are selected at random. This is equivalent to the one-way ANOVA fixed effects model (Shrout & Fleiss, 1979).

## Acknowledgments

This work was supported by the following grants: T32NS047987 (DMS), R01MH118370 (CG), NSF CAREER2048066 (CG).

Also, DMS would like to thank The Therapeutic Cognitive Neuroscience Fund.

## Data and code availability

Midnight Scan Club data is publicly available (https://openneuro.org/datasets/ds000224).

Code related to the analysis in this paper is located at https://github.com/DijunQuant/communication_public/.

## Footnotes

1 Although, in the present work, we do not attempt to address the directionality in the local coupling.

2 In the following text, we may omit the word change when we refer to *communication change* for convenience.

3 Although, as discussed in *Introduction*, correlation is only one of many possible ways to estimate functional connectivity.

4 Although, ICC is theoretically bounded by 0 and 1, in practice it is possible to obtain negative, unbounded values due to sampling issues (Bartko, 1976). For example, when the sample size is small, the estimated within-group variance may exceed the variance of the whole population. It is often recommended that negative values be cut off. Although the threshold can affect the ICC statistics, our conclusion does not change whether or not we apply the cutoffs.

## Notes

### Competing Interest Statement

The authors have declared no competing interest.

