## Supplements for "From Correlation to Communication: disentangling hidden factors from functional connectivity changes"

S1

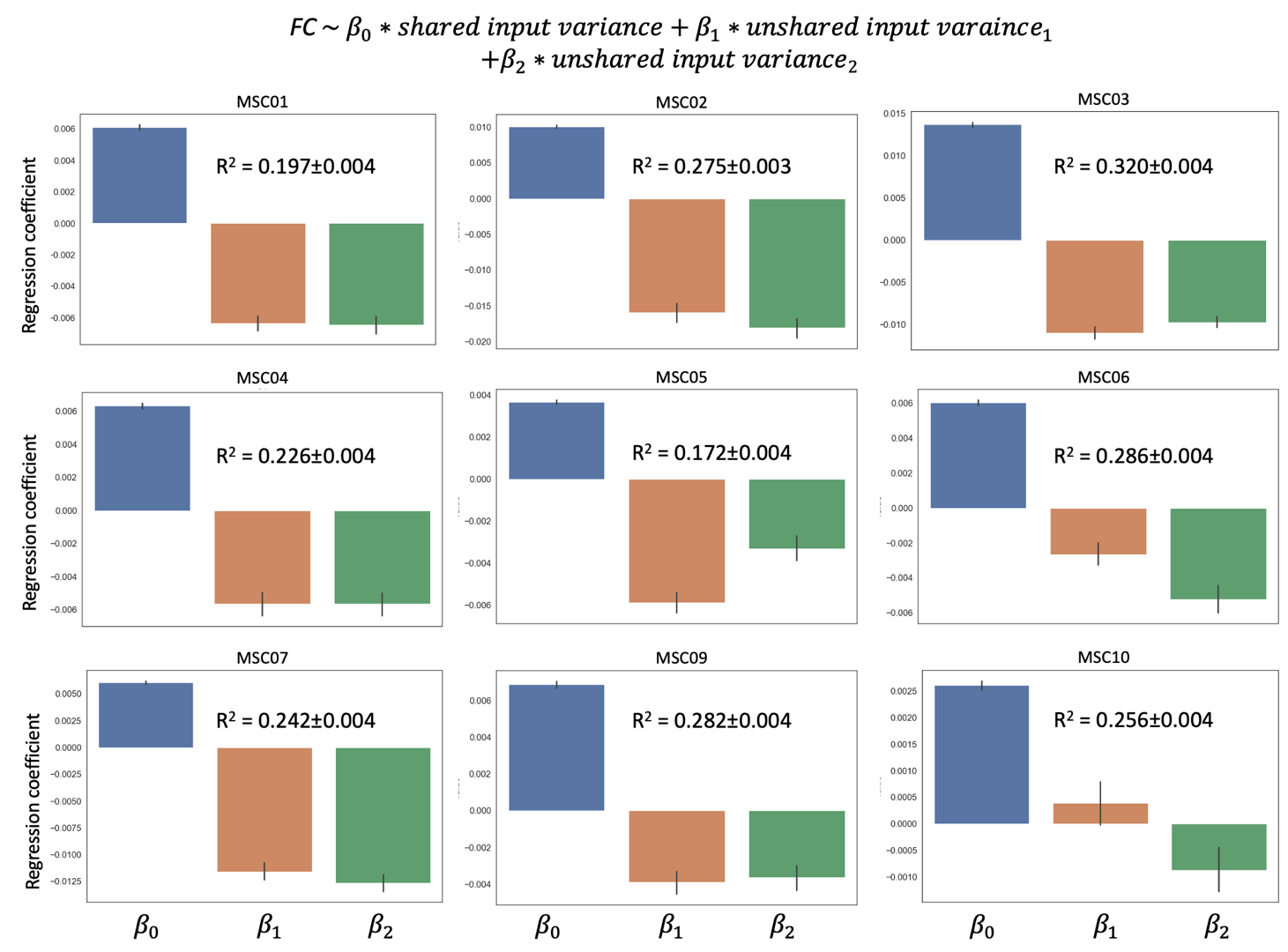

Figure S1. Distribution of regression coefficients of FC against unshared and unshared input variance in the MSC fMRI data. For each edge, a linear regression is fit to 10 resting state scans. Error bar shows the standard error across edges. Each graph shows the result from one participant in the MSC dataset.

S2

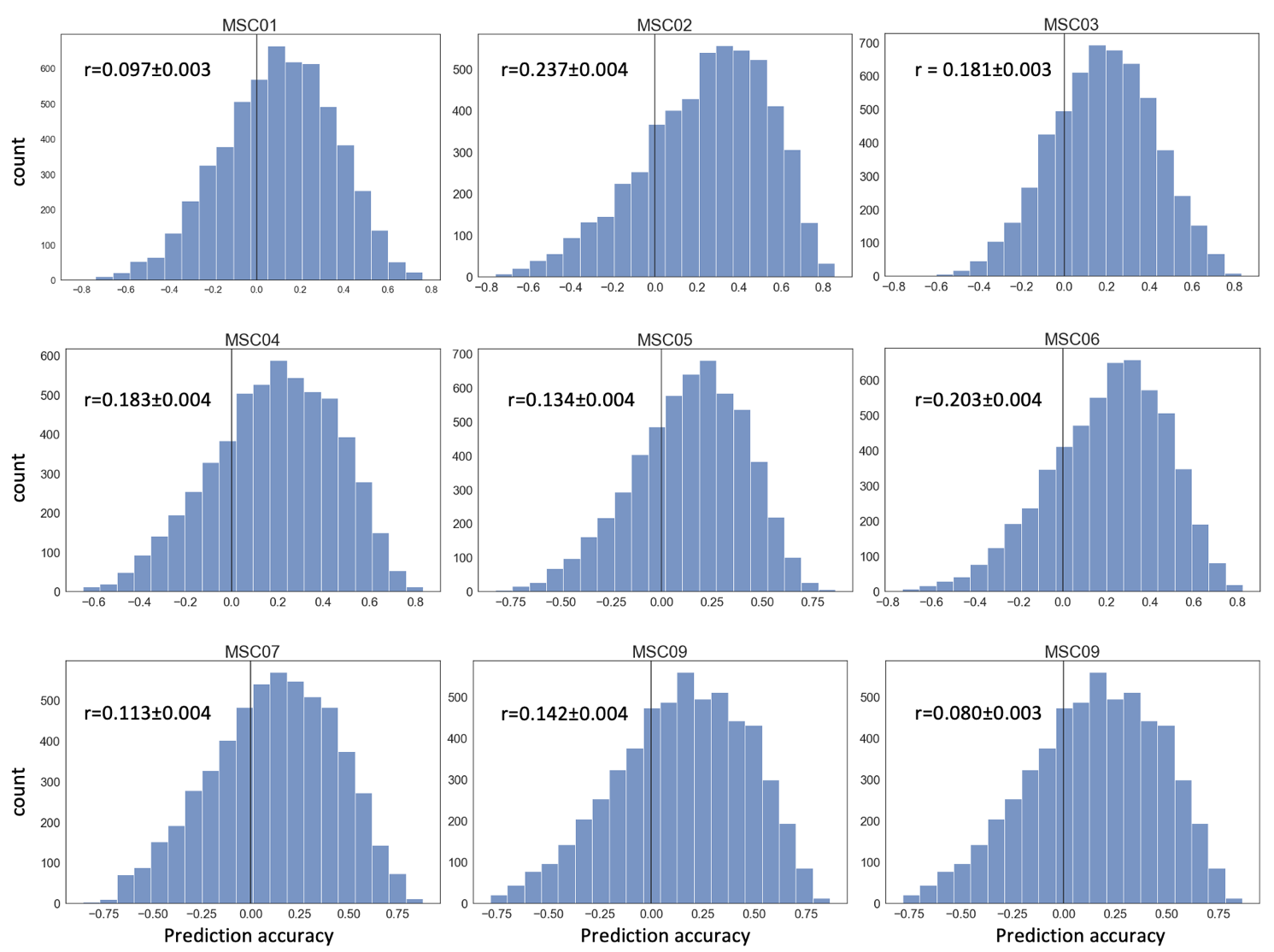

Figure S2. Histogram (number of edges) of prediction accuracies for task-induced FC in each participant from the Midnight Scan Club. For each edge, accuracy is measured by correlation coefficients between actual FC changes and those predicted by task-evoked input variance changes across 30 task sessions (10 sessions for memory tasks, 10 for motor task, and 10 for mixed task). Each graph depicts the result for a single participant from MSC dataset. Mean and standard error are shown on each subplot. For sub09, motor task was excluded due to excess movement.

S3

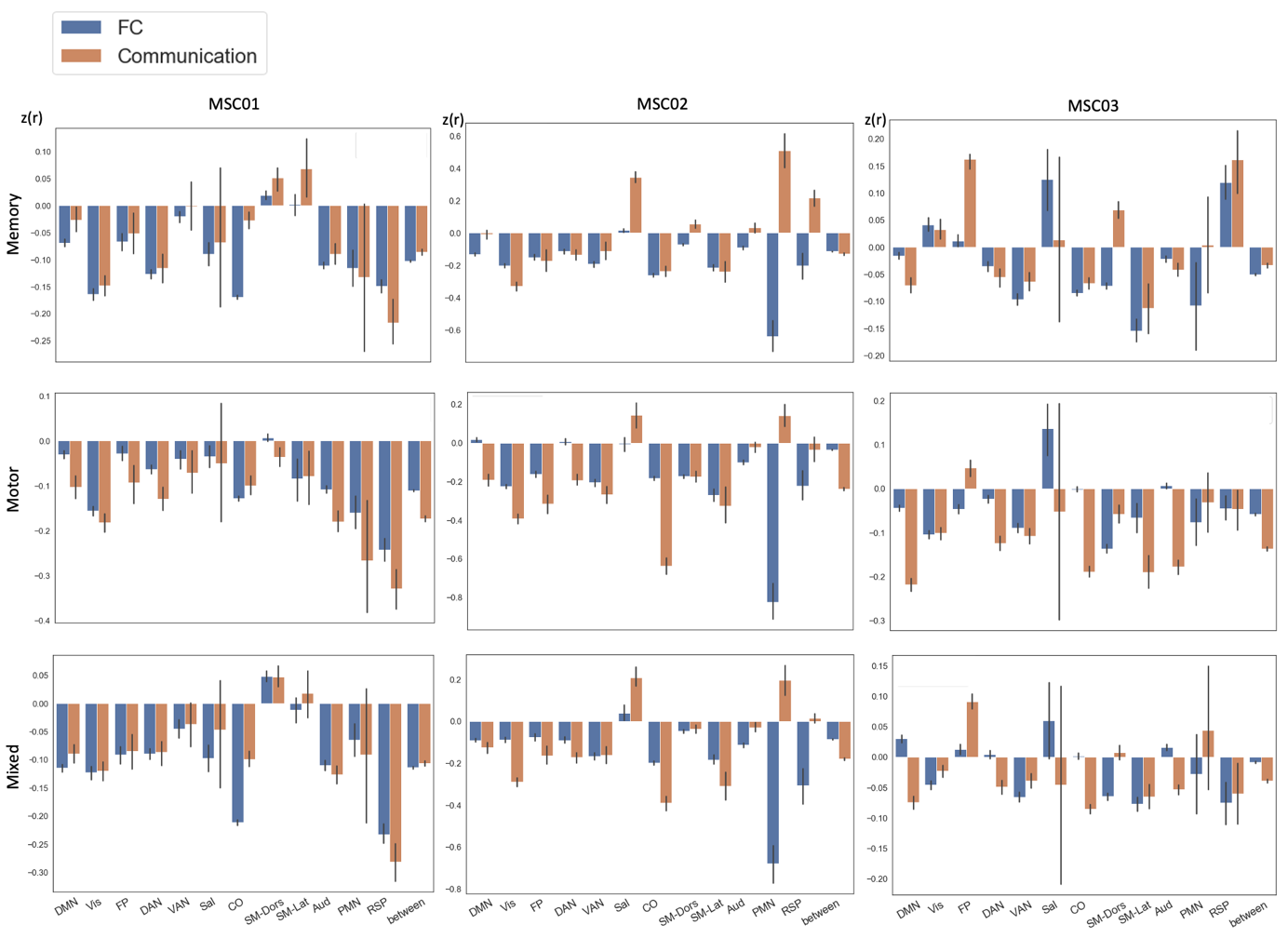

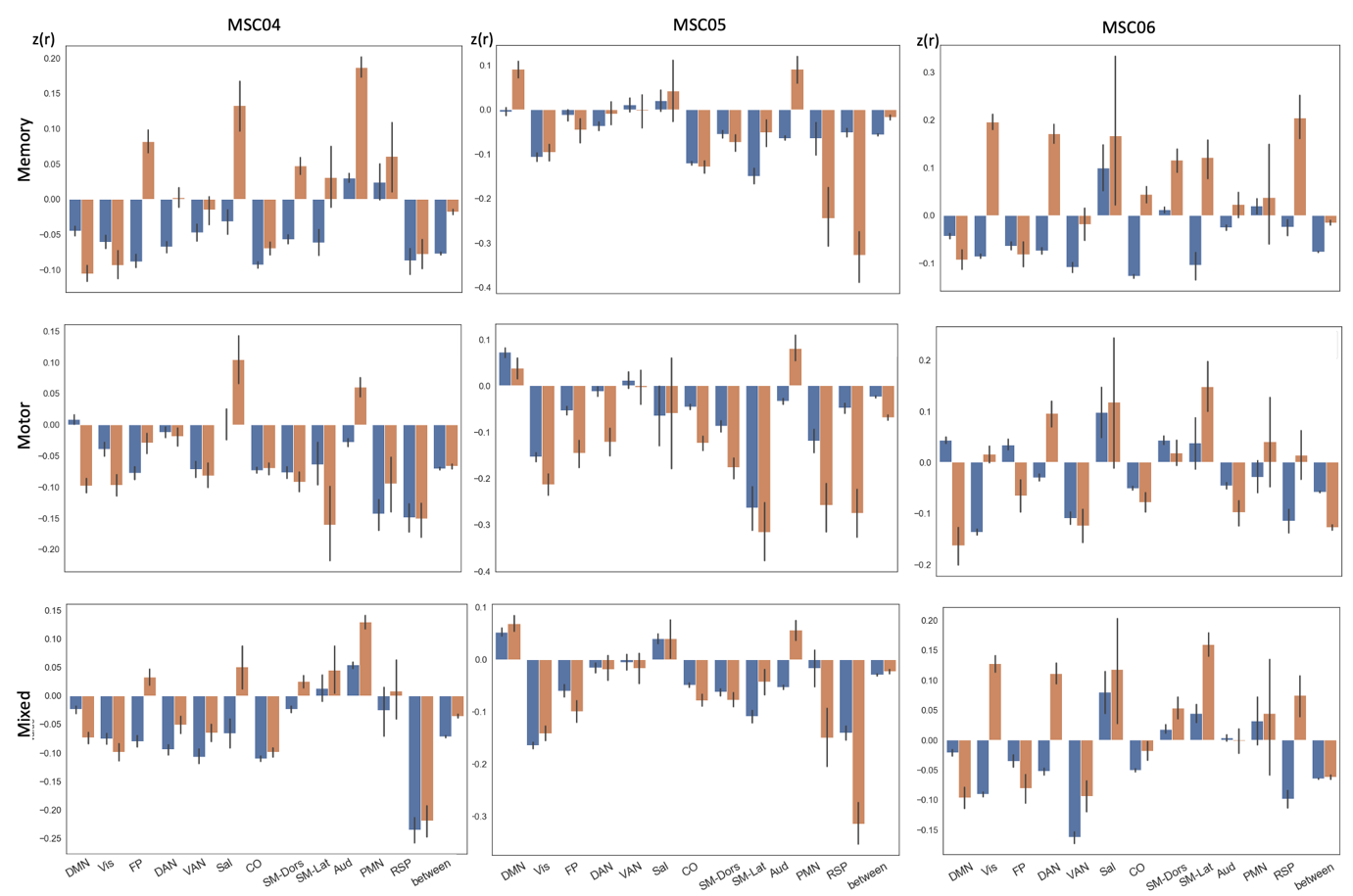

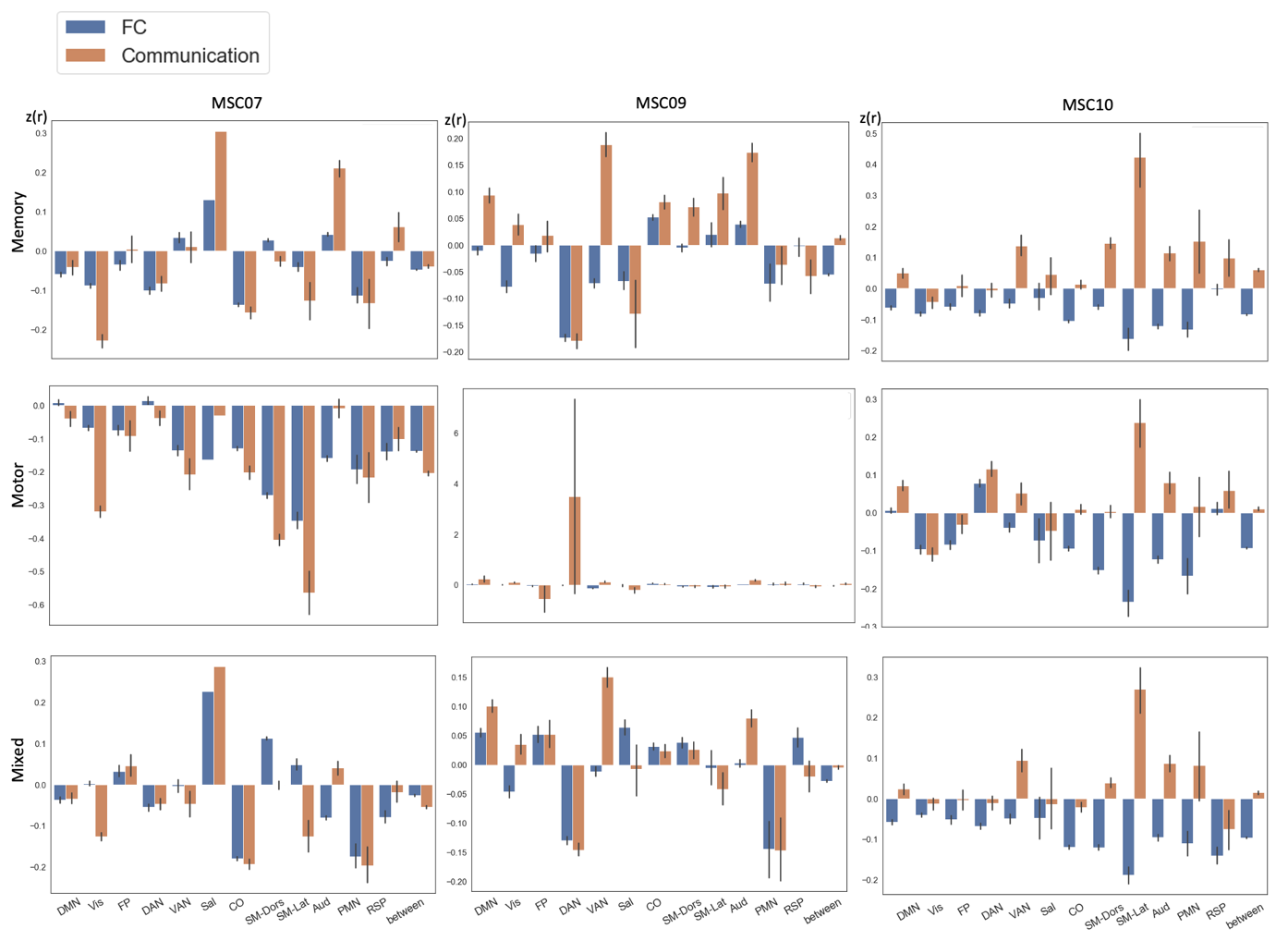

Figure S3. Task-induced FC and communication changes for all edges, grouped by network affiliation during memory task (Memory), motor task (Motor), and mixed task (Mixed) for participants in the Midnight Scan Club. Error bar indicates a standard error across edges. (A) Each set of three graphs in a column depict the result from one individual from the MSC dataset.

S4

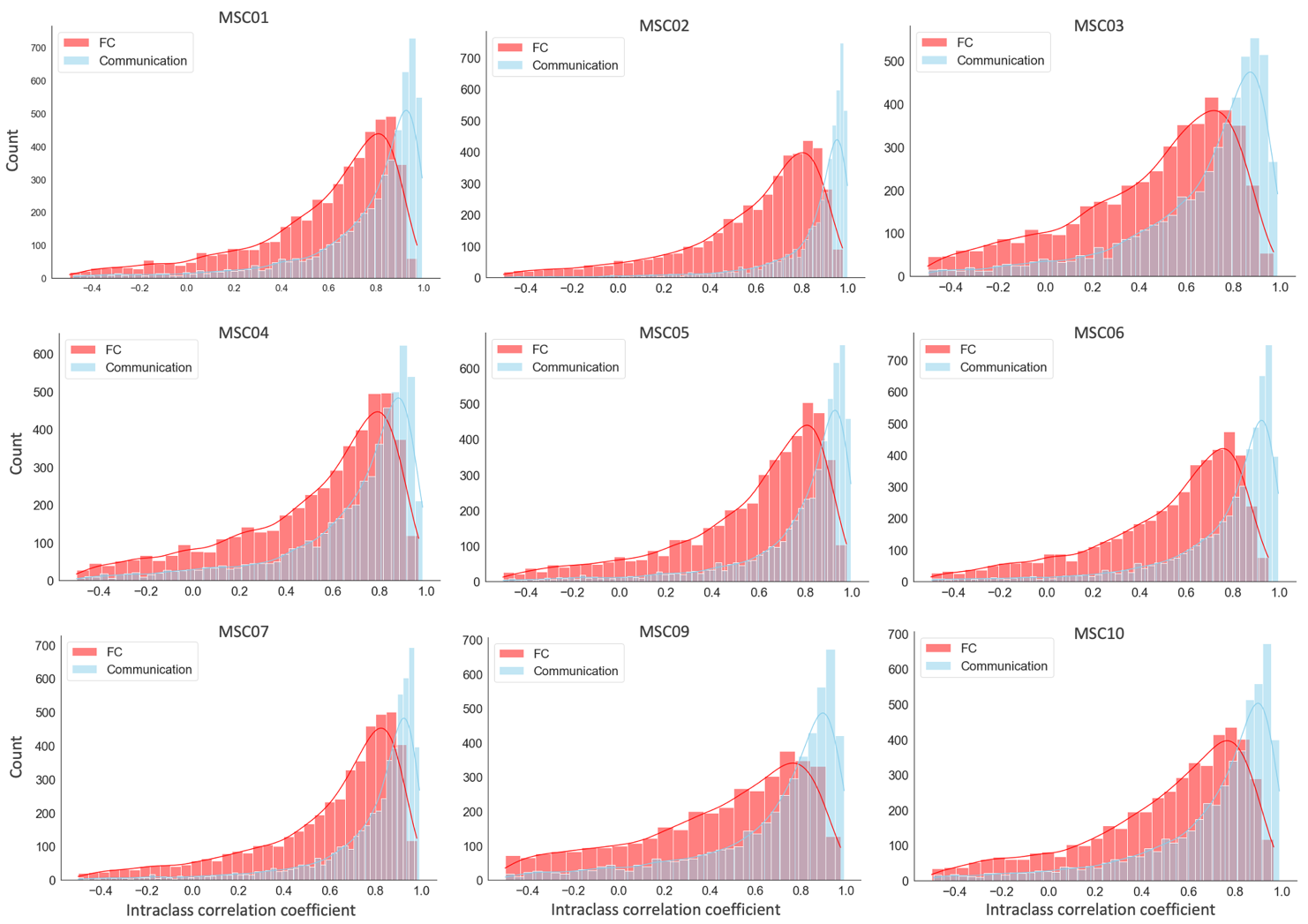

Figure S4. Distribution of Intraclass correlation coefficients (ICC) across edges each participant in MSC dataset. ICC between task types was computed for each edge across 40 scan conditions.

S5

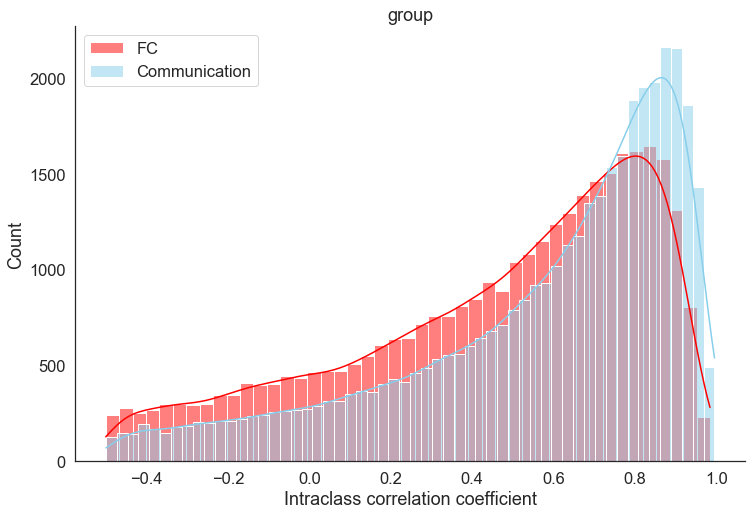

Figure S5. Distribution of Intraclass correlation coefficients (ICC) for all participants. ICC between task types was computed for each edge across 30 task scan session (excluding the rest state).

S6

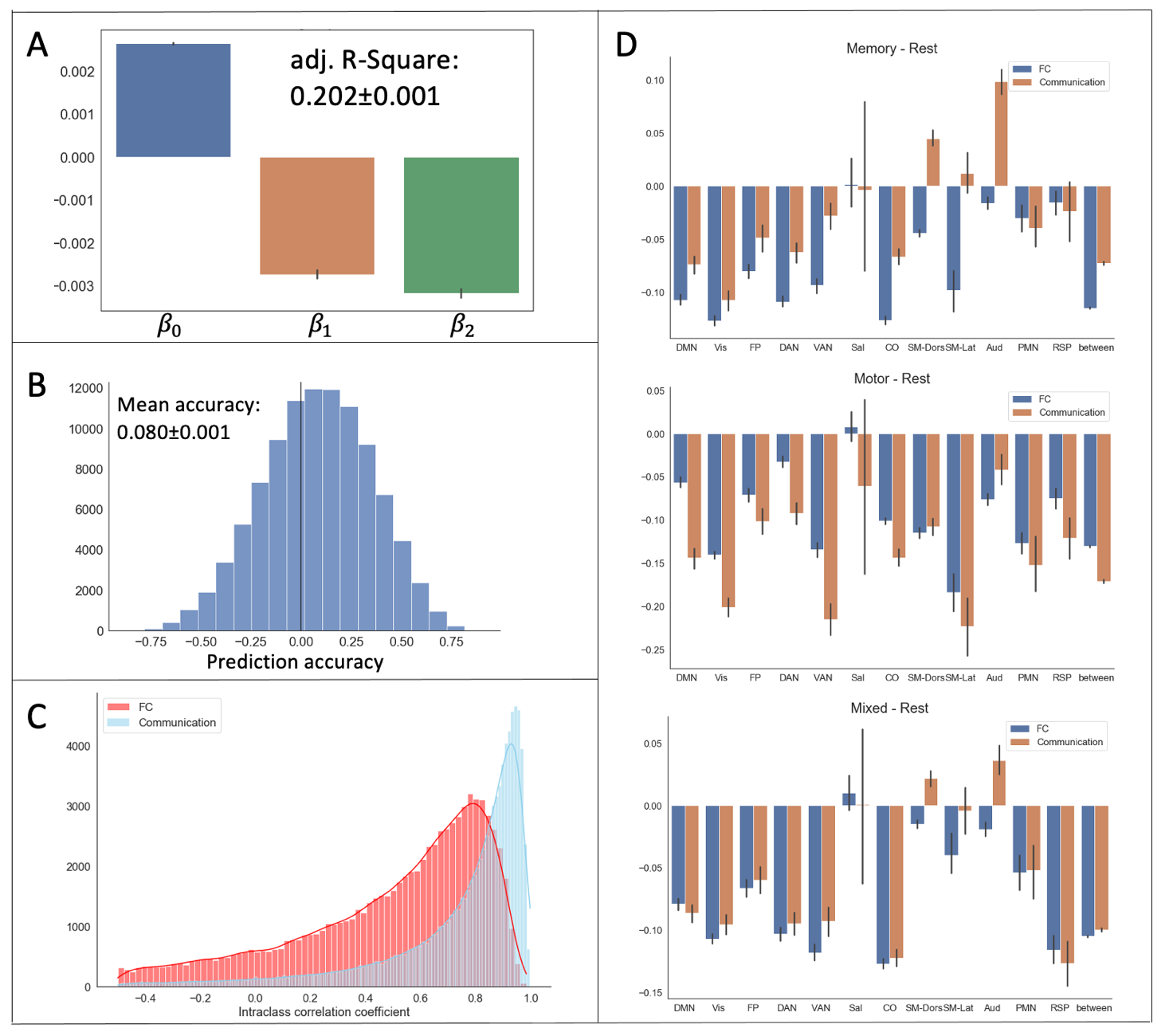

*Figure S6. Key findings replicated using a sparsity level of 10% for selecting network edges. Group data is presented for the same set of analysis in the main text. (A) Distribution of regression coefficients of FC against the unshared and* *unshared input variance during the resting state. (B) Distribution of prediction accuracies, measured by correlation coefficients, for all edges. (C) Distribution of Intraclass correlation coefficients (ICC) for discriminating four task types. (D) Task-induced FC and communication change for all edges grouped by network affiliation during the memory task, motor task and mixed task.*
